# LRP::FLAG reduces phosphorylated tau levels in Alzheimer’s Disease

**DOI:** 10.1101/2020.01.14.905661

**Authors:** Katelyn Cuttler, Monique J. Bignoux, Tyrone C. Otgaar, Stephanie Chigumba, Eloise Ferreira, Stefan F.T. Weiss

## Abstract

Alzheimer’s disease (AD) is characterized by amyloid beta (Aβ) plaque and neurofibrillary tangle formation, respectively. Neurofibrillary tangles form as a result of the intracellular accumulation of hyperphosphorylated tau. Telomerase activity and levels of the human reverse transcriptase (hTERT) subunit of telomerase are significantly decreased in AD. Recently it has been demonstrated that the 37kDa/67kDa laminin receptor (LRP/LR) interacts with telomerase and is implicated in Aβ pathology. Here we show that LRP/LR co-localizes with tau in the perinuclear cell compartment and FRET confirmed a direct interaction between LRP/LR and tau in HEK-293 cells. Overexpression of LRP::FLAG in HEK-293 and SH-SY5Y cells decreased total and phosphorylated tau levels with a concomitant decrease in PrP^c^ levels, a tauopathy-related protein. Additionally, LRP::FLAG overexpression resulted in increased hTERT levels. These data indicate for the first time a role of LRP/LR in tauopathy of Alzheimer’s Disease and recommend LRP::FLAG as a potential alternative therapeutic tool for Alzheimer’s Disease treatment through rescuing cells from Aβ induced cytotoxicity and, as shown in this report, decreased phosphorylated tau levels.

## Introduction

Alzheimer’s disease (AD) which affects approximately 50 million people worldwide [1]. It is the most prevalent form of dementia and incidence is rapidly increasing. Since AD mainly affects the elderly, it is an increasing concern due to increased global life expectancy. In addition, only longterm palliative care is available for AD patients and therefore, AD is a significant socio-economic burden. Indeed, the cost of specialized care for AD amounted to $1 trillion dollars worldwide in 2018 [1]. Thus, there is a great need to develop effective disease-modifying treatments for AD.

There are two key neuropathological hallmarks in AD, namely amyloid plaques formed from the extracellular aggregation of amyloid beta (Aβ) and neurofibrillary tangles formed from the intracellular accumulation of hyperphosphorylated tau [2]. While Aβ is considered by some to be the candidate etiological agent of AD [3, 4], hyperphosphorylated tau is equally important in the progression of AD. Neurofibrillary tangles accumulate in specific brain regions involved in memory and block the microtubular transport system, which harms synaptic communication [2], while amyloid plaques disrupt neuronal functions, resulting in apoptosis. Taken together, these agents are able to damage many areas of the brain, ultimately resulting in death. Therefore, it is vital to develop treatments that target hyperphosphorylated tau as well as Aβ.

Tau is a microtubule-associated protein that normally functions to promote the assembly of tubulin into microtubules and to bind to and stabilize assembled microtubules [5]. Since these microtubules are situated along cytoskeletal tracks, they mediate vital cell processes, including the transport of nutrients and neurotransmitters. In healthy cells, the level of tau phosphorylation is highly modulated and regulates microtubule plasticity [6], motor activity and axonal transport [7], as well as neurite outgrowth [8]. In AD, however, this process is dysregulated, and tau is hyperphosphorylated and dissociates from microtubules. This hyperphosphorylated tau then diffuses into the cytoplasm and associates to form neurofibrillary tangles [9]. This causes cytoskeletal tracks to disintegrate, thereby inhibiting the vital cell processes they mediate, which ultimately leads to cell death [9]. Overall, hyperphosphorylation of tau negatively impacts synaptic health, plasma membrane cell signalling and the protection of DNA from cell stressors [9], all of which may contribute to the progression of AD. Indeed, studies have shown that the degree of dementia is correlated with the number of neurofibrillary tangles present within the brain tissue [10–14].

Furthermore, hyperphosphorylation of tau can be caused by highly neurotoxic Aβ fibrils which activate the tau kinase, glycogen synthase kinase 3 (GSK3) [5, 15]. Therefore, tau hyperphosphorylation is dependent on the extracellular aggregation of Aβ [16]. Thus, the aggregation of Aβ is necessary in order for both the Aβ pathology and tauopathy of AD. In order to do this, Aβ needs to interact with lipid membranes and cellular receptors [17]. One such receptor is the 37 kDa/ 67 kDa Laminin Receptor Precursor/ High Affinity Laminin Receptor (LRP/LR) [18], also known as LamR1, RPSA and p40 [19]. LRP/LR is a multifunctional receptor mainly found in the lipid raft regions of plasma membranes, as well as in the nucleus [20] and cytoplasm [21]. LRP/LR is implicated in numerous diseases, including cancer, prion disorders, ageing and AD. LRP/LR is upregulated in multiple metastatic cancers causing increased metastasis, increased angiogenesis and the impediment of apoptosis [22–34]. Furthermore, LRP/LR is also involved in both cancer and ageing via a functional relationship with telomerase [35–37].

In AD, LRP/LR, PrP^c^ and Aβ have been shown to function together in order for the disease to progress. LRP/LR acts as a receptor for Aβ_42_ internalization, co-localizes with amyloid precursor protein (APP), γ-secretase and β-secretase, and interacts directly with the presenilin 1 (PS1) subunit of γ-secretase during the amyloidogenic processing of APP [18,38–40]. Moreover, blockage or downregulation of LRP/LR with the anti-LRP/LR specific antibody, IgG1-iS18, or short hairpin RNAs (shRNAs), respectively, has successfully impeded Aβ shedding and rescued cells from Aβ-induced cytotoxicity [38, 39]. Furthermore, it was found that the IgG1-iS18 antibody-mediated rescuing of cells from cytotoxicity is dependent on PrP^c^ due to the interaction between PrP^c^ and Aβ42 [41]. Recently, an *in vivo* study, whereby B6SJL-Tg6799 AD transgenic mice were injected with the IgG1-iS18 antibody, confirmed the *in vitro* findings whereby a decrease in amyloid plaque formation and Aβ42 levels with a concomitant increase in both shortterm and learning memory was observed [42].

Another protein recently found to be associated with AD is telomerase. Telomerase is a multisubunit ribonucleoprotein which functions to add TTAGGG repeats to the ends of chromosomes [43]. In highly proliferative cells, the addition of these repeats protects telomeres from erosion [43]. There are two essential components to the human telomerase enzyme, namely hTERC, the RNA component, and hTERT, the reverse transcriptase [44]. Telomerase activity is thus important in cellular senescence and is implicated in both ageing and cancer where it has opposing effects [45, 46]. Importantly, cellular ageing occurs as a result of limited telomerase activity. The resultant telomere erosion causes cells to enter replicative senescence where they are no longer able to divide [46]. This process is supposed to act as a defense mechanism against cancer formation [47–49], however, it can lead to the development of age-related disorders. Furthermore, hTERT has numerous extra-telomeric functions. It is known to function as a transcription factor in several pathways, such as the Wnt-β-catenin pathway and NF-κB pathway, which are both involved in cellular proliferation [50]. Additionally, hTERT is able to migrate to mitochondria when cells are under oxidative stress, in order to convey protection against reactive oxygen species (ROS) [51, 52]. Furthermore, hTERT plays a role in the replication and repair of mitochondrial DNA (mtDNA), thereby protecting cells from mtDNA damage [53]. Since AD is considered an age-related disorder, telomerase has also been implicated in AD. Indeed, telomerase activity is known to be inhibited by Aβ and cells affected by an aggregation of Aβ have critically shortened telomeres [54, 55]. Additionally, hTERT is able to persist in neurons where it may play a neuroprotective role against hyperphosphorylated tau and the resultant neurofibrillary tangles [56]. Indeed, the absence of hTERT further contributes to neurodegeneration as it can no longer protect mitochondria from oxidative stress [56]. Furthermore, overexpressing hTERT is known to protect cells against Aβ-induced apoptosis [57]. Therefore, an upregulation of telomerase, and specifically hTERT, may be a potential treatment strategy for AD as it may be able to protect neurons from both Aβ and tau-induced damage [55, 56].

Recently, Otgaar *et al*., [36] found that an overexpression of LRP::FLAG increased both hTERT levels and telomerase activity with a concomitant reduction in senescent markers. Furthermore, Ferreira *et al*., [42] observed an increase in mTERT in mice treated with the anti-LRP/LR specific antibody, IgG1-iS18 with a concomitant decrease in Aβ peptides and amyloid plaques. In addition, an increase in telomerase activity and hTERT expression was observed with a concomitant reduction in Aβ shedding and intracellular Aβ levels in cells overexpressing LRP::FLAG [58]. Furthermore, this increase in hTERT rescued cells from Aβ-mediated cytotoxicity [58]. Therefore, LRP/LR has a role in the upregulation of telomerase activity and hTERT expression. Since both LRP/LR and telomerase are known to play a role in the Aβ facet of AD, it was hypothesized that they could also play a role in tauopathy. Thus, this study aimed to determine if LRP/LR has a relationship with tau and whether overexpression of LRP::FLAG has an effect on tau and tauopathy-related proteins.

## Results and Discussion

LRP/LR is essential for Aβ mediated cytotoxicity during the development and progression of AD. However, it is not known whether LRP/LR has an effect on the other neuropathological agent of AD, the hyperphosphorylated tau. Therefore, this study demonstrates that LRP/LR interacts directly with tau and that LRP::FLAG overexpression decreases the levels of total and phosphorylated forms of tau, as well as PrP^c^. Concurrently, the overexpression of LRP::FLAG also increased the levels of hTERT and phosphorylated hTERT.

### LRP/LR interacts directly with tau

In order to determine whether LRP/LR interacts with tau, confocal microscopy with Airyscan™ was first utilized to confirm whether these two proteins co-localized in HEK-293 cells (Figure 1A). Figure 1A d-f represent a merge of tau (red) with LRP/LR (green), and areas of yellow indicate spatial overlap. Since both LRP/LR and tau are reportedly found in the cytoplasm and the nucleus, our results confirm this and further indicate that these proteins co-localise in those cell compartments [20, 21]. Since we observed the co-localisation of LRP/LR and tau, we wanted to investigate whether there was an interaction between these proteins. FRET analysis was employed in HEK-293 cells and confirmed that LRP and tau interact. FRET occurs between fluorescently labelled proteins if they are within 10 nm of each other (Förster, 1948), resulting in the fluorescence shift which we observed in this study (Figure 1B f). Therefore, not only do LRP/LR and tau co-localise, our results indicate that these proteins directly interact, as they exist within 10 nm of each other.

**Figure 1.**
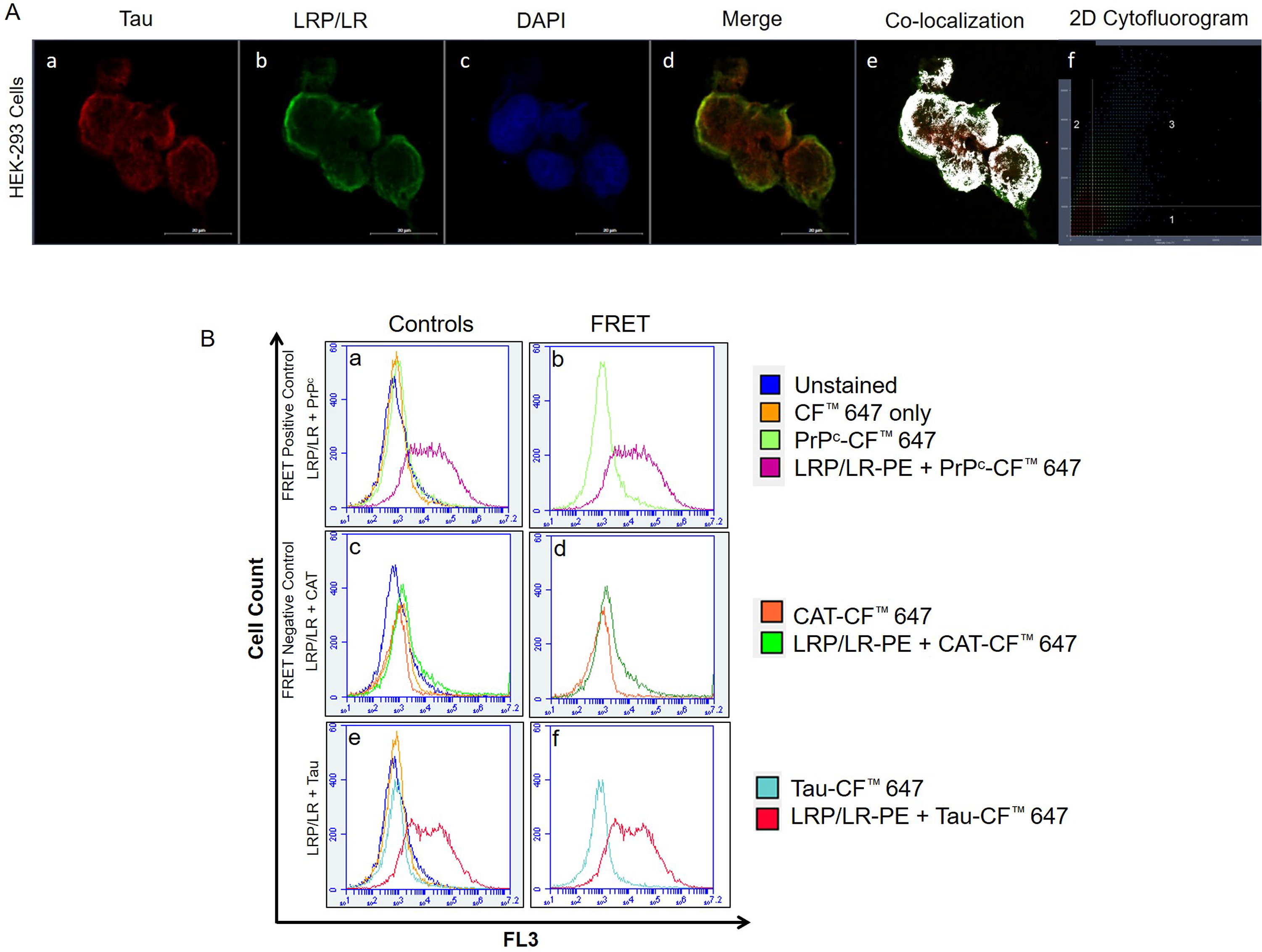
LRP/LR and tau co-localize and interact in HEK-293 cells. (A) Image showing the co-localization between tau (red) (a) and LRP/LR (green) (b). Nuclei are stained with DAPI (blue) (c). Co-localization occurs between LRP/LR and tau in the perinuclear compartment, represented by yellow fluorescence in the merged image (d), as white areas (e) and as fluorescence in the third quadrant of the 2D cytofluorogram (f). All images were taken at 630x magnification with the addition of Airyscan^™^. Scale bars represent 20 μm. (B) Flow cytometric analysis of FRET between intracellular LRP/LR and tau in HEK-293 cells shows a shift in fluorescence which is indicative of a direct interaction. The fluorescence intensity of the unstained cells (dark blue histogram) was superimposed with that of cells labelled with the CF^™^ 647 secondary antibody only (yellow histogram) as well as with cells in which the proteins of interest; PrP^c^, CAT and tau, were labelled with CF^™^ 647 (a, c, e). (b) Co-labelling of LRP/LR-PE and PrPc-CF^™^ 647 (positive control) (pink histogram). (d) Co-labelling of LRP/LR-PE and CAT-CF^™^ 647 (negative control) (green histogram). (f) Co-labelling of LRP/LR-PE and Tau-CF^™^ 647 (red histogram). Each panel is a representative image. n = 3 biological repeats.

Since, LRP/LR and tau were shown to interact directly (Figure 1B), it is thus possible that LRP/LR is involved in the main function of tau; to assemble and maintain microtubules in healthy cells. Indeed, cytoplasmic LRP is known to tightly associate with tubulin structures and may serve to tether ribosomes to microtubules in order to assist in protein synthesis [59, 60]. Therefore, it is possible that cytoplasmic LRP could also bind to tau peptides in order to assist in stabilizing microtubules during their assembly and normal functioning. In AD, however, this interaction could be misappropriated to worsen the destabilizing effects of tau hyperphosphorylation. Cytoplasmic LRP may no longer be able to stabilize microtubules and this could contribute to the disintegration of cytoskeletal tracks. Importantly, though, strategies targeting LRP/LR could make use of this interaction for therapeutic intervention in AD by counteracting the microtubule destabilizing effects of tau hyperphosphorylation.

### LRP::FLAG overexpression decreases total and phosphorylated tau levels

Since a direct interaction was confirmed between LRP/LR and tau, we wanted to investigate whether LRP/LR plays a role in tauopathy. HEK-293 and SH-SY5Y cells were stably transfected with the pCIneo-moLRP-FLAG plasmid [61] to induce an overexpression of LRP::FLAG. Western blotting confirmed the transfection of both HEK-293 and SH-SY5Y cells (Figure 2A; C; Figure S2). After confirming transfection of both cell lines, western blotting was utilized to determine the effect of LRP::FLAG overexpression on both total and phosphorylated tau levels (Figure 2A; C; Figure S3-S5). A significant decrease in total tau of 45% and 35% was observed in transfected HEK-293 (Figure 2B) and transfected SH-SY5Y cells (Figure 2D), respectively. Since it is not possible to investigate hyperphosphorylation of tau *in vitro*, two antibodies were used which target tau at two different phosphorylation sites (pS404 and pT231). In transfected HEK-293 cells there was a significant 68% reduction in tau phosphorylated at S404 and a significant 98% reduction in tau phosphorylated at T231 (Figure 2B). In transfected SH-SY5Y cells, a significant 63% and 92% reduction was observed, respectively (Figure 2D). Since these are two of the major culprits of tauopathy, the decreased phosphorylation at these two sites together may indicate a possible decrease in hyperphosphorylated tau [62–65]. This decrease in phosphorylated tau was also observed in the cytoplasm and the nucleus of both HEK-293 and SH-SY5Y cells (Figure 3) by using confocal microscopy. Therefore the overexpression of LRP::FLAG might decrease the overall hyperphosphorylation of tau.

**Figure 2:**
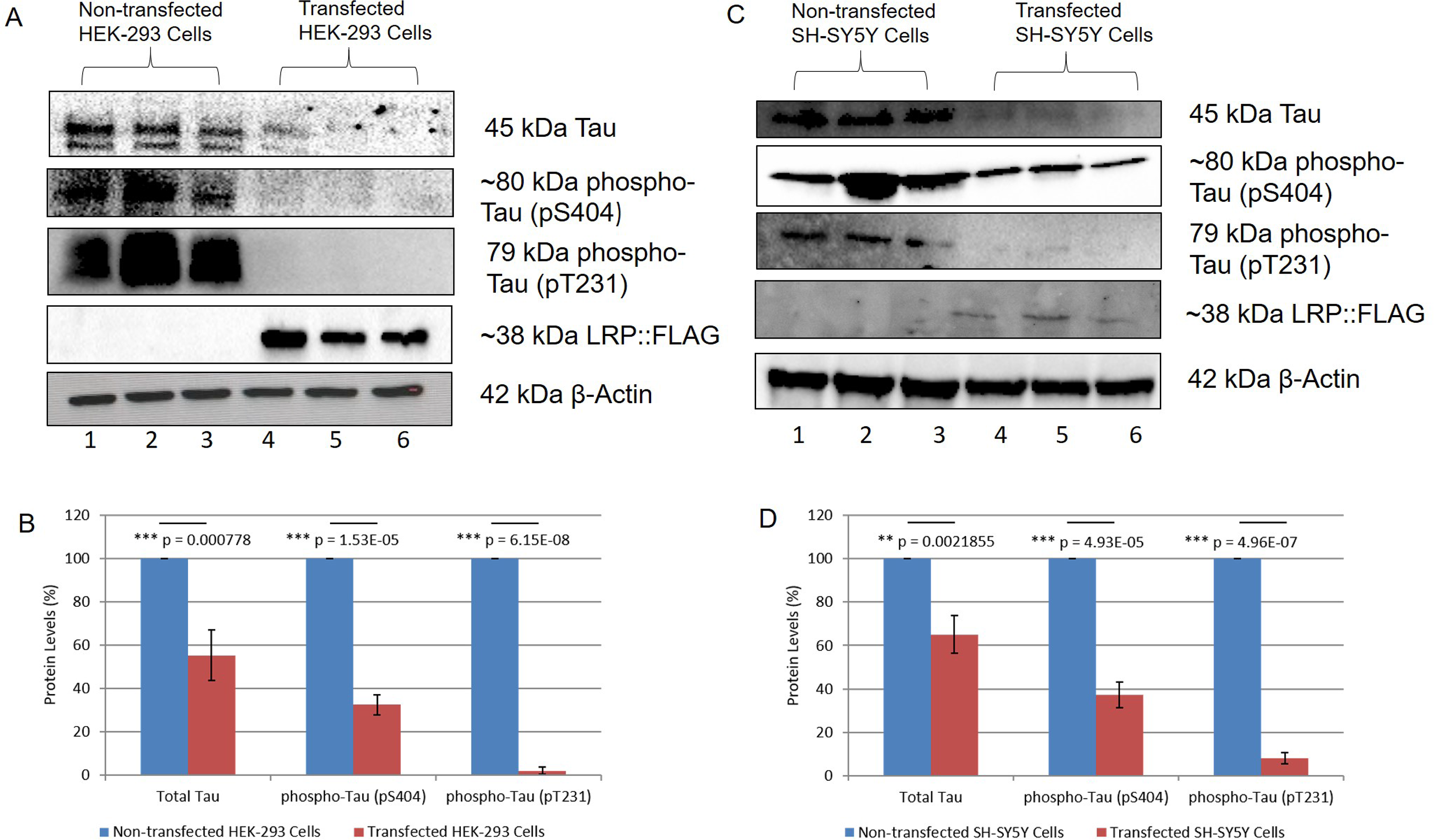
Overexpression of LRP::FLAG in HEK-293 and SH-SY5Y cells decreases levels of total Tau as well as phospho-Tau. A, B) A 45% decrease in total Tau, 68% decrease in phospho-Tau (pS404) and 98% decrease in phospho-Tau (pT231) was observed in transfected HEK-293 cells. C, D) A 35% decrease in total Tau, 63% decrease in phospho-Tau (pS404) and 92% decrease in phospho-Tau (pT231) was observed in transfected SH-SY5Y cells. A, C) HEK-293 and SH-SY5Y cells were confirmed to be overexpressing LRP::FLAG after stable transfection with the pCIneo-moLRP-FLAG plasmid. LRP::FLAG was detected in transfected HEK-293 cells only (A). LRP::FLAG was detected in transfected SHSY-5Y cells only (C). A, C) β-Actin was used as a loading control. Densitometric analysis performed was relative to the non-transfected HEK-293 (B) and SH-SY5Y (C) cells which were set to 100%. Error bars represent standard deviation, n=3 biological repeats. *p: 0.05, **p: 0.01, ***p: 0.001; Student’s *t*-test.

**Figure 3:**
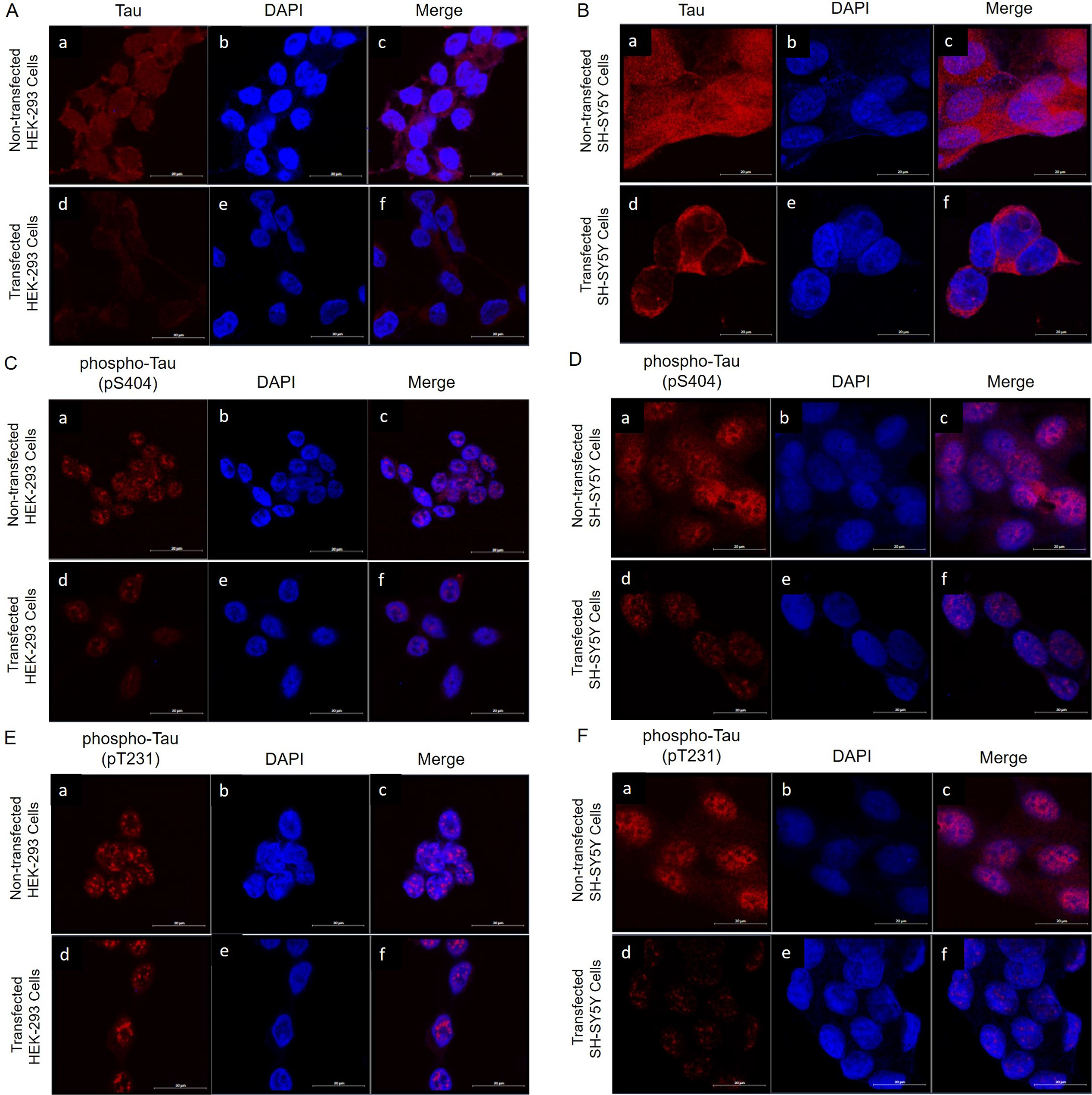
Intracellular expression of tau and phospho-Tau in transfected and non-transfected HEK-293 and SH-SY5Y cells. b, e) Nuclei are stained with DAPI (blue). A, B) Expression of total tau decreases in transfected HEK-293 and transfected SH-SY5Y cells. Endogenous levels of total tau (red) (a). Levels of total tau (red) after transfection with pCIneo-moLRP-FLAG plasmid (d). Merge of total tau with DAPI staining (c, f). C, D) Expression of phospho-Tau (pS404) is decreased in transfected HEK-293 and transfected SH-SY5Y cells. Endogenous levels of phospho-Tau (pS404) (red) (a). Levels of phospho-Tau (pS404) (red) after transfection with pCIneo-moLRP-FLAG plasmid (d). Merge of phospho-Tau (pS404) with DAPI staining (c, f). E, F) Expression of phospho-Tau (pT231) is decreased in transfected HEK-293 and transfected SH-SY5Y cells. Endogenous levels of phospho-Tau (pT231) (red) (a). Levels of phospho-Tau (pT231) (red) after transfection with pCIneo-moLRP-FLAG plasmid (d). Merge of phospho-Tau (pT231) with DAPI staining (c, f). All images were taken at 630x magnification with the addition of Airyscan^™^. Scale bars represent 20 μm.

Hyperphosphorylation of tau is a major problem in AD where it forms tangles that disrupt the internal structure of neuronal cells by de-stabilizing the cytoskeletal tracks [9]. Thus the significant decrease observed in both tau phosphorylation sites (pS404 and pT231) may result in a reduced amount of aggregation of hyperphosphorylated tau, resulting in less damage to neuronal cells. [62–65] Since this aggregation is known to cause disintegration of cytoskeletal tracks, a decrease in hyperphosphorylated tau may hinder cytoskeleton disintegration and improve vital cell processes, such as axonal transport, thereby permitting neurons to retain normal functioning in AD patients [9]. In particular, the phosphorylation of tau at S404 and T231, which was investigated in this study, is known to negatively affect the binding of the tau protein to the microtubules, thereby de-stabilizing them and inhibiting cytoskeletal signalling and transport [65, 66]. Indeed, T231 mutant N18 neuroblastoma cells abolished phosphorylation of tau by the serine/threonine kinase GSK3, indicating that phosphorylation at T231 plays an important role in the hyperphosphorylation of tau [63]. Therefore, a decrease in phosphorylation at these sites may prevent the dissociation of tau and promote tubulin polymerization and assembly into microtubules [63, 66]. Thus, the structural integrity of the cells would be maintained, and their signalling and transport capabilities would remain intact, allowing them to better resist the damaging effects of AD. In particular, DNA damage repair proteins can be trafficked to the sites of cytotoxicity in order to rescue the cells [67]. The DNA damage caused by the cytotoxicity present in AD results in neuronal apoptosis via oxidative stress [68, 69]. In addition, hyperphosphorylation of tau is able to enhance Aβ cytotoxicity by sequestering Fyn kinase [9]. Fyn is thus re-distributed to the neuron whereby it can strengthen the excitotoxic signalling of the neurotransmitter glutamate, which is known to enhance the cytotoxicity of Aβ oligomers [9, 70]. Therefore, a decrease in phosphorylated tau and the resultant decrease in neurofibrillary tangles could both lessen cytotoxicity as well as allow DNA repair to occur. Overall, the decrease in phosphorylated tau observed could be acting as a potential mitigator for AD development and progression.

Furthermore, a decrease in total tau levels was observed in both cell lines (Figure 2, Figure 3). Wild-type tau is essential for the functioning of microtubules and cell signalling, therefore a decrease in total tau may be detrimental to the cells. However, we have seen that LRP::FLAG overexpression rescues cells from cytotoxicity after treatment with Aβ42 [58]. Therefore, we suggest that the effect of the decreased tau is minimal, otherwise the cells would enter apoptosis as a result of microtubule disintegration and loss of structural integrity. Indeed, there is a known redundancy between tau and other microtubule-associated proteins [71], such as the redundancy between MAP-1B and tau during axonal growth [72, 73]. Therefore, it is possible that another protein could function in the place of tau in order to maintain the internal structure of the cell. In addition, the decreases in total tau are more finite than those seen in phosphorylated tau. Thus, while overexpression of LRP::FLAG does not specifically target phosphorylated tau, it seems to have a much more notable effect on the phosphorylated form. In this regard, it is known that the interaction between LRP and laminin is able to affect the phosphorylation of ERK, JNK and p38 proteins [74]. Therefore, LRP may be involved in kinase/phosphatase axes and laminin cell signalling. Thus, it is possible that the LRP-laminin interaction could also affect the phosphorylation of tau, especially as LRP/LR was shown to directly interact with tau. Furthermore, LRP/LR and focal adhesion kinase (FAK) have been shown to interact after LRP/LR binds to laminin [75]. Indeed, in neurons treated with Aβ there is also increased phosphorylation of FAK and an increased association between FAK and Fyn kinase in addition to the increased phosphorylation of tau. Taken together, these increases are known to enhance cytotoxicity in AD [76]. Therefore, it is possible that the interaction between LRP and FAK, and the subsequent activation of signalling pathways, such as PI3-kinase/Akt and MAPK [75], may affect the phosphorylation of tau in AD. Therefore, given the potential role of LRP in kinase/phosphatase axes and that LRP and tau directly interact, it is plausible that LRP could likewise affect the phosphorylation of tau.

### LRP::FLAG overexpression increases hTERT and phospho-TERT levels

LRP/LR has also recently been found to interact with the anti-ageing related proteins telomerase and hTERT [36]. In AD, neuronal cells have critically shortened telomeres as an accumulation of Aβ is able to inhibit the activity of telomerase [54, 55]. Furthermore, there is an absence of hTERT in cells affected by hyperphosphorylated tau [56]. Since both LRP/LR and telomerase are known to play a role in the Aβ facet of AD, it was hypothesized that they could also play a role in tauopathy. It is known that overexpression of LRP::FLAG increases hTERT levels in HEK-293 cells [36] and SH-SY5Y cells [58]. It has also been shown that this increase results in an increase in telomerase activity [36,58]. Thus, western blotting was performed to examine the effect of LRP::FLAG overexpression on hTERT levels, together with the active form of hTERT, phospho-TERT (pS824) (Figure 4A; C; Figure S6-S7), as this is indicative of an increase in telomerase activity. A significant increase of 120% and 125% in hTERT levels was observed in transfected HEK-293 (Figure 4B) and SH-SY5Y (Figure 4D) cells, respectively. This is consistent with previous observations [36, 58]. A significant increase in phospho-TERT levels was also observed both cell lines, 112% in transfected HEK-293 cells (Figure 4B) and 96% in transfected SH-SH5Y cells (Figure 4D). It is known that an increase in hTERT induces/ causes a corresponding increase in telomerase activity [36,77,78]. However, in order to be active, hTERT has to be present in the nucleus [79]. Therefore, in order for nuclear localisation of the full length 120kDa hTERT variant to occur, it firstly undergoes phosphorylation at the serine residues 227 and/or 824 by PI3-kinase/AKT signalling [79]. This suggests that the increase in telomerase activity is as a result of not only increased hTERT levels but also hTERT import to the nucleus. Therefore, the increase in both hTERT and phospho-TERT observed in transfected HEK-293 and SH-SY5Y cells is indicative of an increase in telomerase activity as previously observed by Otgaar et al. [36] and Bignoux et al. [58].

**Figure 4.**
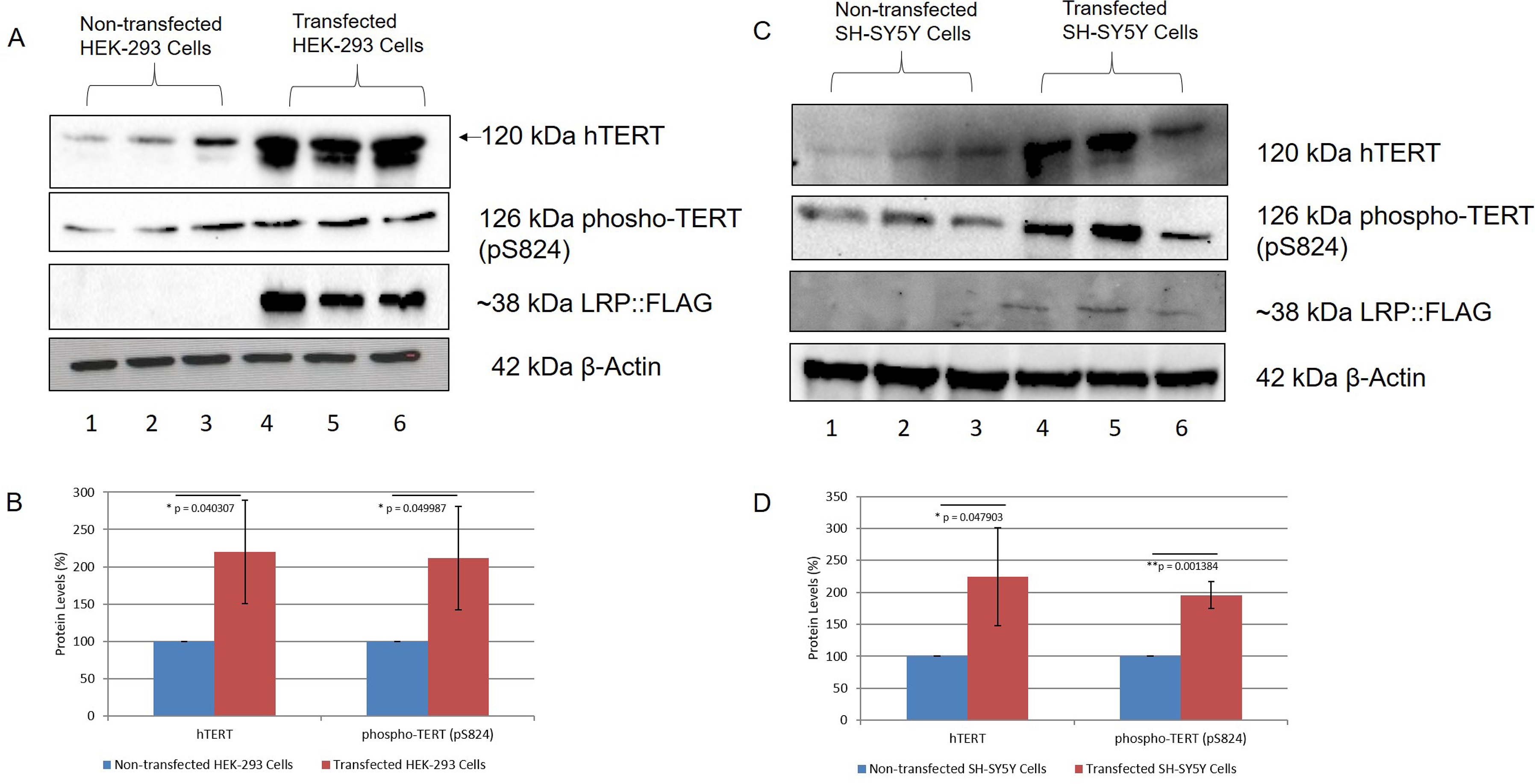
LRP::FLAG overexpression increases hTERT levels in HEK-293 and SH-SY5Y cells and increases phospho-TERT levels in SH-SY5Y cells. A, B) A 120% increase in hTERT and 112% increase in phospho-TERT was observed in transfected HEK-293 cells. C, D) A 125% increase in hTERT and 96% increase in phospho-TERT was observed in transfected SH-SY5Y cells. β-Actin was used as a loading control. Densitometric analysis performed was relative to the non-transfected HEK-293 (B) and SH-SY5Y (D) cells which were set to 100%. Error bars represent standard deviation, n=3 biological repeats. *p: 0.05, **p: 0.01, ***p: 0.001; Student’s *t*-test.

Indeed, by comparing phospho-TERT and hTERT levels in HEK-293 and SH-SY5Y cells, it is clear that not all hTERT is being converted into phospho-TERT. This hTERT may thus be involved in extra-telomeric functions. Of note is the protective function of hTERT against ROS. hTERT is able to migrate to the mitochondria when cells are under oxidative stress, which allows hTERT to interact with mtDNA in order to convey protection [51, 52]. In AD, there is compromised energy production in the brain due to mitochondrial dysfunction [80]. This dysfunction can increase ROS production which has been shown to promote the hyperphosphorylation of tau [81], although it has not yet been determined how this occurs. Conversely, hyperphosphorylation of tau and the formation of neurofibrillary tangles could also impair the movement of mitochondria along microtubules [82]. In neurons, mitochondria are distributed to areas of the axon where metabolic demand is high, such as synapses [83, 84], and active growth cones or branches [85, 86]. As a result, ATP depletion can occur in other areas of axons and dendrites which can worsen synaptic dysfunction [87]. Importantly, in AD, the absence of hTERT has been shown to increase the levels of mitochondrial superoxide in neurons [56] and truncated tau expressed *in vivo* generates increased oxidative stress over time through an accumulation of ROS [88]. Furthermore, pathological tau and hTERT have been observed to be mutually exclusive [56], suggesting that neurons expressing hTERT may be prevented from developing tau pathology. Indeed, it has previously been observed that LRP::FLAG overexpression causes an increase in hTERT in the cytoplasm and nucleus [36, 58] and it is this increase in the cytoplasm and nucleus which could correspond to an increase in the extra-telomeric functions of hTERT. Therefore, in AD, the increase in hTERT observed, particularly in the cytoplasm, could improve mitochondrial function and protect against ROS production. Furthermore, nuclear hTERT is able to regulate inflammatory and proliferative pathways, such as Wnt and NF-κB [50]. Therefore, the increased nuclear hTERT observed could regulate pro-proliferative and anti-inflammatory genes via these pathways which could prevent apoptosis and aid cell survival in AD. Therefore, an increase in hTERT may prevent cell cycle arrest and apoptosis in a number of ways; through increased telomerase activity and telomere elongation, by reducing ROS production thereby improving mitochondrial function, as well as by regulating cell survival pathways. Overall, an increase in hTERT could slow the progression of tauopathy and AD.

### LRP::FLAG overexpression decreases PrP^c^ levels

Although the role of PrP^c^ in tau hyperphosphorylation is poorly understood, it is the primary receptor for Aβ42 and has an important role in Aβ42 pathology [89]. Thus, it was hypothesized that PrP^c^ may also be important for tauopathy.

The effect of LRP::FLAG overexpression on PrP^c^ was therefore investigated using western blotting (Figure 5A, 5C; Figure S8). Endogenous PrP^c^ can exist as three isoforms: nonglycosylated, monoglycosylated and diglycosylated and thus ranges in size from approximately 20 kDa to approximately 35 kDa (Vana and Weiss, 2006). In HEK-293 cells, both the di- and monoglycosylated forms were seen (Figure 5A), while in SH-SY5Y cells only the monoglycosylated form was observed (Figure 5C). It is important to note that PrP^c^ glycoform ratios differ across brain regions and across cell lines, thereby explaining the different banding patterns [90, 91]. Therefore, the differences in total PrP^c^ levels were used for comparison. A significant decrease in total PrP^c^ in both transfected HEK-293 and SH-SY5Y cells of 31% (Figure 5B) and 39% (Figure 5D), respectively, was observed.

**Figure 5:**
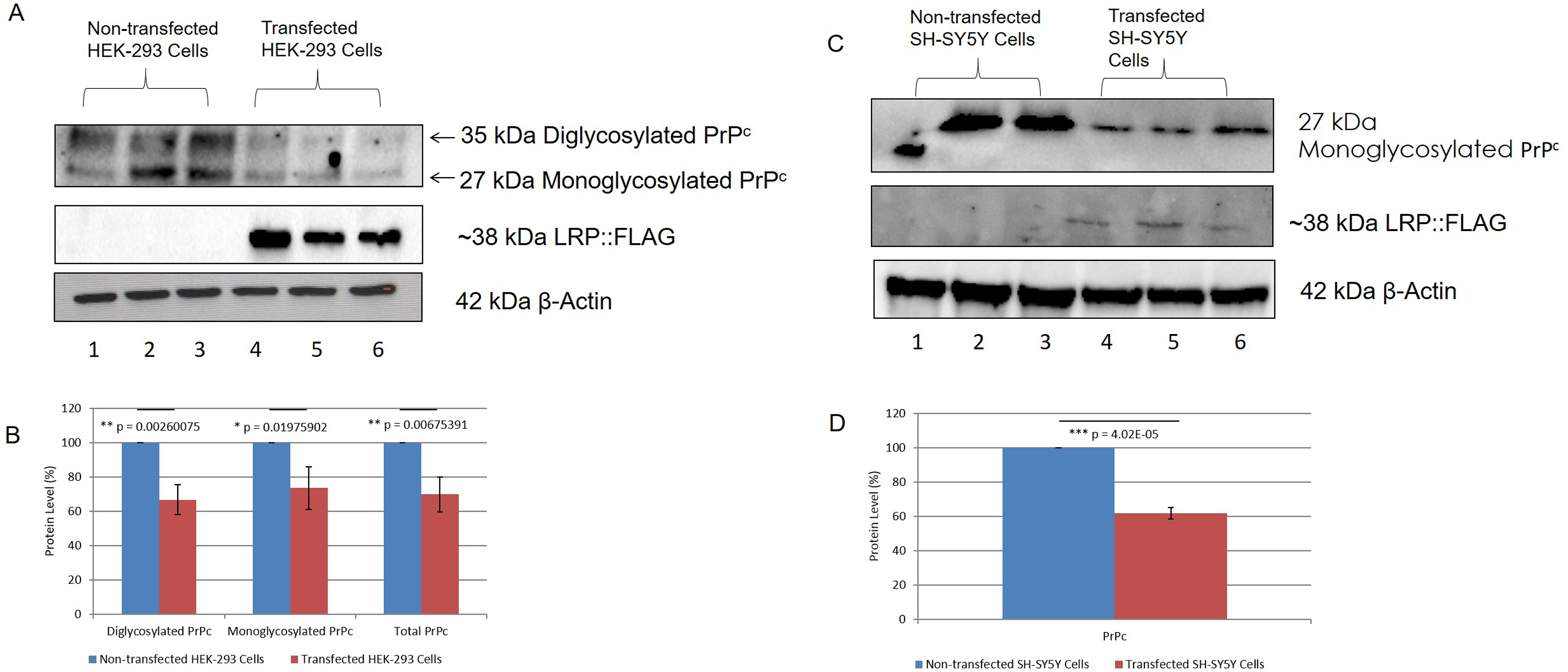
Overexpression of LRP::FLAG in HEK-293 and SH-SY5Y cells decreases PrP^c^ levels. A, B) A 31% decrease in total PrP^c^ (di- and monoglycosylated), 34% decrease in diglycosylated PrP^c^ and 27% decrease in monoglycosylated PrP^c^ was observed in transfected HEK-293 cells. (C, D) A 39% decrease in total PrP^c^ (monoglycosylated) was observed in transfected SH-SY5Y cells. β-Actin was used as a loading control. Densitometric analysis performed was relative to the non-transfected HEK-293 (B) and SH-SY5Y (D) cells which were set to 100%. Error bars represent standard deviation, n=3 biological repeats. *p: 0.05, **p: 0.01, ***p: 0.001; Student’s *t*-test.

It has been suggested that PrP^c^ has a protective role against tau hyperphosphorylation and is able to downregulate tau expression [92, 93]. Contrary to this, we observed a decrease in both PrP^c^ and phosphorylated tau in cells overexpressing LRP::FLAG. However, it has previously been observed that LRP::FLAG overexpression specifically reduced both intracellular and extracellular Aβ42 levels [58]. Indeed, Larson *et al*. [94] identified a signalling cascade between soluble Aβ and PrP^c^ which activates Fyn kinase. This activated Fyn kinase is known to cause synaptic and cognitive impairment in transgenic mice when dysregulated [95, 96]. Furthermore, Larson *et al*. [94] discovered that this PrP^c^-dependent activation of Fyn can also cause the missorting and hyperphosphorylation of tau. However, high levels of Aβ are required for this PrP^c^-mediated Fyn activation and this may only occur during the later stages of AD [94]. Therefore, we postulate that PrP^c^ may switch roles as AD progresses due to the increasing levels of cytotoxic Aβ. Thus, in the current study, the decrease in PrP^c^ observed may be beneficial in preventing the Aβ-PrP^c^ signalling cascade occurring in the later stages of AD. Therefore LRP::FLAG overexpression is able to decrease phosphorylated tau, Aβ42 and PrP^c^ levels. These proteins interact with each other during progression of AD, whereby Aβ42 binds to PrP^c^ causing cytotoxicity and increased Aβ42 shedding, which results in hyperphosphorylation of tau and the formation of neurofibrillary tangles. Furthermore, as the diagnosis and knowledge of AD improves, it may become easier to determine when PrP^c^ becomes detrimental and thus a potential target of treatment. Overall, we believe that the decrease in PrP^c^ caused by LRP::FLAG overexpression could be beneficial; however more research would need to be done to better understand the role of PrP^c^ in tauopathy.

### Conclusion

In conclusion, LRP/LR and tau share the same sub-cellular locations and directly interact with each other. LRP::FLAG overexpression caused a decrease in phosphorylated tau and PrP^c^ levels. Taken together with previous results showing that LRP::FLAG overexpression decreases Aβ42 levels as well as rescuing cells from Aβ42-mediated cytotoxicity [58], this study shows that the overexpression of LRP::FLAG is able to affect each of the major proteins in the Aβ42-PrP^c^-hyperphosphorylated tau signalling cascade and thereby decrease the cytotoxic effects observed in AD. Furthermore, hTERT levels increased after LRP::FLAG overexpression and thus could provide protection against the cytotoxicity caused by tau hyperphosphorylation. LRP::FLAG might act as a potential alternative therapeutic tool for Alzheimer’s Disease treatment through a rescuing mechanism of A-ß mediated cytotoxicity and a decrease of phosphorylated tau levels which might be mediated by increased hTERT and decreased PrP^c^ levels, respectively.

## Methods

### Cell Culture

HEK293 and SH-SY5Y cells were cultured in DMEM with high glucose (4.5 g/l) and 4 mM L-Glutamine (Hyclone): Ham’s F12 with 1 mM L-Glutamine (Hyclone) in a 1:1 ratio. In addition, the media was supplemented with 15% FBS (Hyclone) and 2% penicillin/streptomycin (Biowest). Cells were maintained at 37 °C in a humidified incubator.

### Confocal Microscopy with the addition of Airyscan^™^

Cells were seeded onto microscope coverslips within a 6-well plate and incubated overnight to reach 50-70% confluency. All further steps were performed with gentle agitation. Cells were fixed with 4% formaldehyde (VWR) in PBS for 20 minutes at room temperature. Cells were then washed 3 times in 1X PBS, after which they were permeabilized with 0.25% Triton X-100 (Sigma Aldrich) in PBS for 20 minutes. Thereafter, cells were washed 3 times in 1X PBS and blocked in 0.5% BSA (Amresco) in PBS for 20 minutes. Cells were then incubated in primary antibody (1:200 in 0.5% BSA in PBS) and incubated overnight at 4 °C. Subsequently, the coverslips were washed 3 times with 0.5% BSA in PBS and then incubated with the appropriate secondary antibody (1:500 in 0.5 % BSA in PBS) for an hour in the dark, at room temperature. The primary and secondary antibodies that were used are listed in Table 1. The cells were then counterstained with 0.1 μg/ml DAPI (Sigma Aldrich) for 5 minutes in the dark for nuclear staining. Coverslips were then washed 3 times with PBS. Coverslips were then mounted onto clean microscope slides using Fluoromount™ Aqueous Mounting Medium (Sigma Aldrich) and incubated at room temperature for 1.5 hours in order to allow the mounting medium to set. Controls were prepared as above, using secondary antibodies only. All images were acquired using the Zeiss LSM 780 confocal microscope at 630x magnification, with the addition of Airyscan™ to obtain resolution of 140 nm. Images were analyzed using Zen 2010 imaging software v2.1.

**Table 1:** Antibodies used for confocal microscopy.

| Target Protein | Primary Antibody | Secondary Antibody |
| --- | --- | --- |
| LRP/LR | Human anti-LRP/LR (IgG1-iS18) | Anti-human IgG-FITC (Sigma Aldrich: F9512) |
| Total Tau (Tau-5) | Mouse anti-Tau (Tau-5) (EMD: MAB361) | Anti-mouse IgG-CF™ 647 (Sigma Aldrich: SAB4600182) |
| Phospho-Tau (pS404) | Rabbit anti-p-Tau (S404) (Cell Signalling: 20194S) | Anti-rabbit IgG-CF™ 647 (Sigma Aldrich: SAB4600184) |
| PHF-Tau (pT231) | Mouse anti-Phospho-Tau (Thr231) (AT180) (Invitrogen: MN1040) | Anti-mouse IgG-CF™ 647 (Sigma Aldrich: SAB4600182) |

### Flow Cytometric Analysis of Försters Resonance Energy Transfer (FRET)

FRET is a very sensitive technique that is used to assess protein interactions *in vitro*. In this experiment the donor/ acceptor pair PE/ APC was used to immunolabel the proteins of interest. The presence of FRET was examined using the FL3 filter set of the Accuri C6 Flow Cytometer (BD Biosciences). CF^™^ 647, which is an APC contemporary, is excited by the 650 nm neon/ helium laser and emits at 660 nm. Within the FL3 channel, excitation is achieved with the 488 nm argon laser and emission is detected at 660 nm. CF^™^ 647 is not excited by the 488 nm laser and therefore does not exhibit fluorescence within this channel. However, if CF^™^ 647 is in close proximity to PE, it may be indirectly excited via FRET resulting in enhanced fluorescence emission in FL3.

The experimental design was based off of the FRET analysis performed by Da Costa Dias *et al*. [18] with a few alterations. Briefly, after a 3 hour incubation in serum-free media, HEK-293 cells were detached and harvested via centrifugation at 150 xg for 10 minutes. Since tau is an intracellular protein, following a 20 minute fixation with 4% formaldehyde at 4 °C, the cells were permeabilized by incubating them in 0.1 % Triton X-100 in PBS at 4 °C for 20 minutes. Thereafter, the cells were blocked in 0.5% BSA in PBS for 10 minutes. Cells were centrifuged at 2700 xg for 10 minutes after each step in order to remove the supernatant. The cells were then incubated in primary antibody solutions for 2 hours, washed thrice with PBS and incubated in secondary antibody solutions for 2 hours. Primary and secondary antibodies were used at a concentration of 1:100 in 0.05% BSA in PBS. The primary and secondary antibodies used are listed in Table 2. Cells were washed in PBS three times prior to analysis. Wash steps were performed by centrifuging cells at 2700 xg for 5 minutes and removing the supernatant. Three biological repeats were performed, and 15 000 cells were analyzed per sample. PrP^c^ was used as a positive control, as there is a direct interaction between LRP/LR and PrP^c^ [97], and chloramphenical acetyl transferase (CAT) was used as a negative control, as LRP/LR has been shown not to interact with it [38].

**Table 2:** Antibodies used for Försters Resonance Energy Transfer (FRET)

| Target Protein | Primary Antibody | Secondary Antibody |
| --- | --- | --- |
| LRP/LR | Human anti-LRP/LR (IgG1-iS18) | Anti-human IgG-PE<br>(Abcam: ab7006) |
| Total Tau (Tau-5) | Mouse anti-Tau (Tau-5)<br>(EMD: MAB361) | Anti-mouse IgG-CF<sup>TM</sup> 647<br>(Sigma Aldrich: SAB4600182) |
| Prion Protein (PrP<sup>c</sup>) | Mouse anti-Prion Protein<br>(Sigma Aldrich: P0110) | Anti-mouse IgG-CF<sup>TM</sup> 647<br>(Sigma Aldrich: SAB4600182) |
| CAT | Rabbit anti-CAT<br>(Sigma Aldrich: C9336) | Anti-rabbit IgG-CF<sup>TM</sup> 647<br>(Sigma Aldrich: SAB4600184) |

### Stable Overexpression of LRP::FLAG

In order to determine the effect of LRP::FLAG overexpression on tau levels, cells were transfected with the plasmid, pCIneo-moLRP-FLAG [61]. Cells were cultured until 40-50% confluency was achieved. The cells were then transfected using the Xfect™ Transfection protocol (Takara). The transfected cells were incubated for 48 hours. Thereafter, cell media was replenished, and the cells were treated with 600-800 ng/ml Geneticin, as a selective treatment. Subsequently, cells were cultured in 200-400 ng/ml of Geneticin to ensure the maintenance of LRP::FLAG expression. Stable transfection was achieved after six weeks of Geneticin treatment.

### Western Blotting

Western blotting was used to confirm expression of LRP::FLAG as well as to detect total protein levels of tau, phospho-tau (pS404 and pT231), hTERT, phospho-TERT and PrP^c^, posttransfection. β-actin was used as the loading control. Briefly, cell lysates were prepared using 1X RIPA buffer and protein levels quantified using a BCA assay. Proteins were resolved on a 12% SDS-PAGE gel for approximately 1 hour at 150 V in 1X electrophoresis buffer (25mM Tris, 192 mM glycine and 0.1% SDS). A total of 30 μg (LRP::FLAG) and 60 μg (Tau, phospho-Tau (pS404), phospho-Tau (pT231), hTERT, phospho-TERT (pS824) and PrP^c^) protein was resolved. Thereafter, the separated proteins were transferred onto a polyvinylidene fluoride (PVDF) membrane using a semi-dry transfer apparatus at 300 mV for 45-50 minutes in 1X transfer buffer (20% methanol in 25 mM Tris and 19.2 mM glycine). The proteins were fixed to the membrane in 0.4% formaldehyde for 30 minutes. Membranes were blocked in 3% BSA in PBS for 1 hour. The blots were incubated overnight with the appropriate primary antibody, diluted in 3% BSA in PBS, with gentle shaking at 4 °C. Subsequently, the membranes were washed in 0.1% Tween 20 in PBS (PBST). The membranes were then incubated with the appropriate secondary antibody, diluted in 3% BSA in PBS, for 1 hour in the dark and, thereafter, washed with PBST. Membranes were incubated with Thermo Scientific chemiluminescent substrate and imaged using the Bio-Rad GelDoc XR Imager. Densitometric analysis was performed using Image Lab 5.1 software (Bio-Rad). Primary and secondary antibodies are listed in Table 3.

**Table 3:** Antibodies used for western blotting.

| Target Protein | Primary Antibody | Dilution | Secondary Antibody | Dilution |
| --- | --- | --- | --- | --- |
| β-Actin | Mouse anti-β-Actin Peroxidase (Sigma Aldrich: A3854) | 1:10000 | - | - |
| LRP::FLAG | Mouse anti-FLAG<sup>®</sup> M2 (Sigma Aldrich: F-3165) | 1:6666 | Anti-mouse IgG-HRP (Abcam: ab97023) | 1:10000 |
| LRP::FLAG | Mouse anti-FLAG<sup>®</sup> M2-Cy3<sup>™</sup> (Sigma Aldrich: A-9594) | 1:5000 | - | - |
| Total Tau (Tau-5) | Mouse anti-Tau (Tau-5) (EMD: MAB361) | 1:500 | Anti-mouse IgG-HRP (Abcam: ab97023) | 1:3333 |
| Phospho-Tau (pS404) | Rabbit anti-p-Tau (S404) (Cell Signalling: 20194S) | 1:1000 | Anti-rabbit IgG-HRP (Cell Signalling: 7074S) | 1:3333 |
| PHF-Tau (pT231) | Mouse anti-Phospho-Tau (Thr231) (AT180) (Invitrogen: MN1040) | 1:500 | Anti-mouse IgG-HRP (Abcam: ab97023) | 1:3333 |
| hTERT | Mouse anti-hTERT | 1:2500 | Anti-mouse IgG-HRP (Abcam: ab97023) | 1:3333 |
|  | (Santa Cruz Biotechnology: sc-77511) |  |  |  |
| Phospho-hTERT (pS824) | Rabbit anti-phospho-Telomerase (pSer<sup>824</sup>) (Sigma Aldrich: SAB4504295) | 1:1000 | Anti-rabbit IgG-HRP (Cell Signalling: 7074S) | 1:3333 |
| Prion Protein (PrP<sup>c</sup>) | Mouse anti-Prion Protein (Sigma Aldrich: P0110) | 1:5000 | Anti-mouse IgG-HRP (Abcam: ab97023) | 1:3333 |

### Statistical Analysis

All statistical analysis of the data obtained was carried out using Microsoft Excel 2016 (Microsoft Corporation). All experiments were performed with three biological repeats and error bars represent standard deviation. The two-tailed Student’s *t*-test was performed at a 95% confidence interval. Values of *p < 0.05 were considered significant, while values of **p < 0.01 and ***p < 0.001 were considered very significant.

## Acknowledgements

This work is based upon research supported by the National Research Foundation (NRF), the Republic of South Africa (RSA), grant numbers 105832, 99061, 118851, awarded to SFTW. Any opinions, findings and conclusions or recommendations expressed in this material are those of the author(s), and therefore, the National Research Foundation does not accept any liability in this regard thereto.

## Author Contributions

KC performed all experiments, analysed the data and wrote the manuscript. MJB and SC assisted with experimental aspects. MJB, TCO and SC assisted with preparation and editing of the manuscript. SFTW and EF conceptualized the study and edited the manuscript.

## Supplementary Material

**Figure S1:**
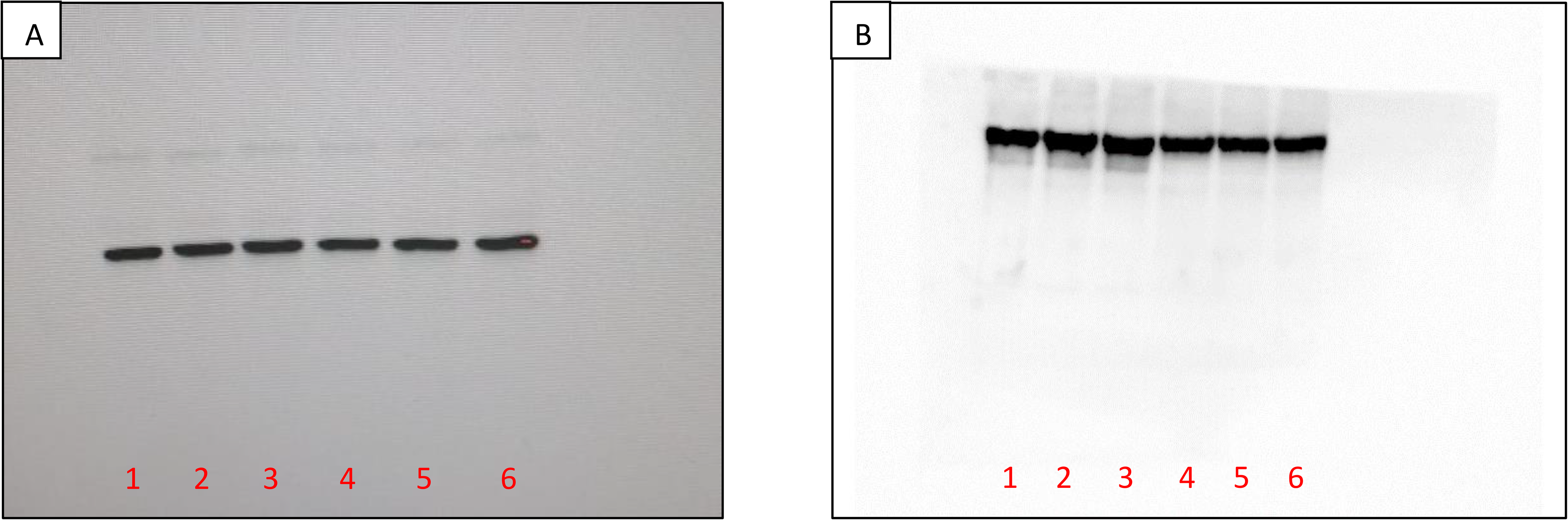
Full β-Actin western blots for HEK-293 cells (A) and SH-SY5Y cells (B). Lanes 1-3: non-transfected cells; lanes 4-6: transfected cells.

**Figure S2:**
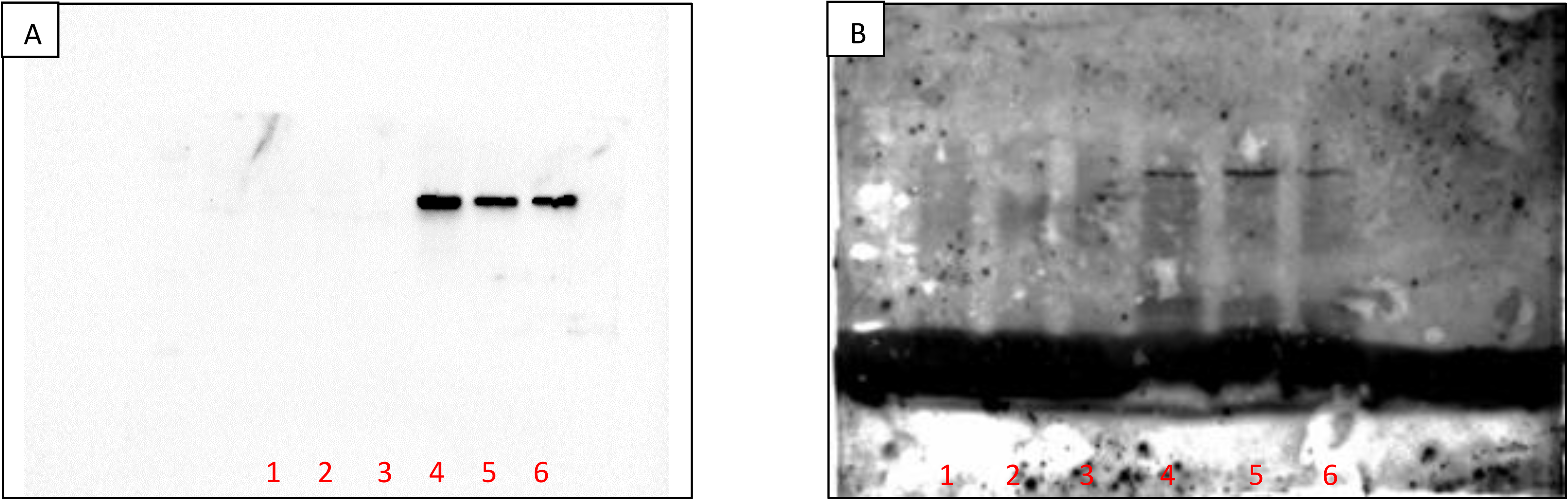
Full FLAG western blots for HEK-293 cells (A) and SH-SY5Y cells (B). Lanes 1-3: non-transfected cells; lanes 4-6: transfected cells.

**Figure S3:**
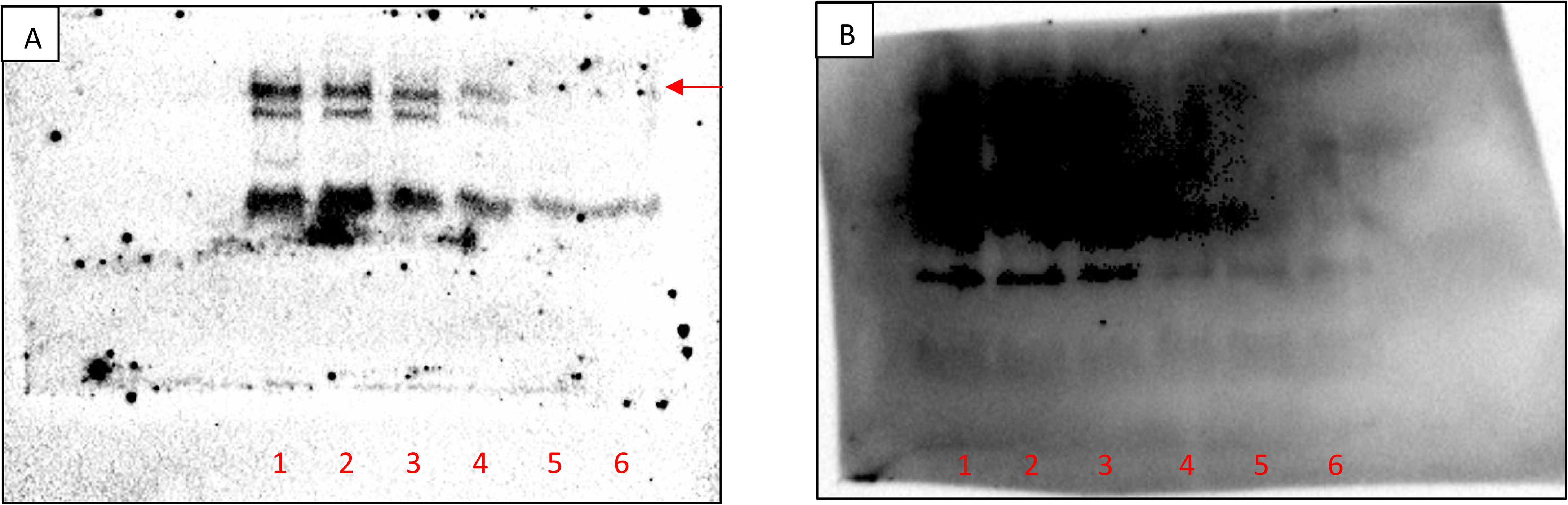
Full total tau western blots for HEK-293 cells (A) and SH-SY5Y cells (B). Lanes 1-3: non-transfected cells; lanes 4-6: transfected cells.

**Figure S4:**
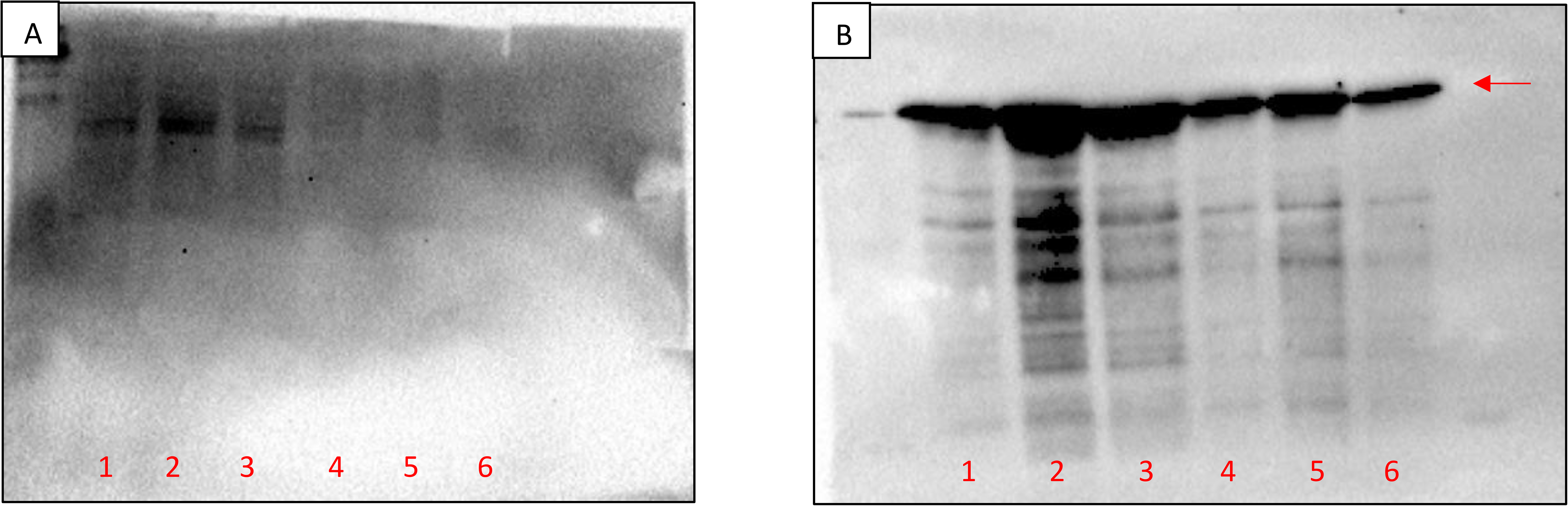
Full phosphorylated tau (pS404) western blots for HEK-293 cells (A) and SH-SY5Y cells (B). Lanes 1-3: non-transfected cells; lanes 4-6: transfected cells.

**Figure S5:**
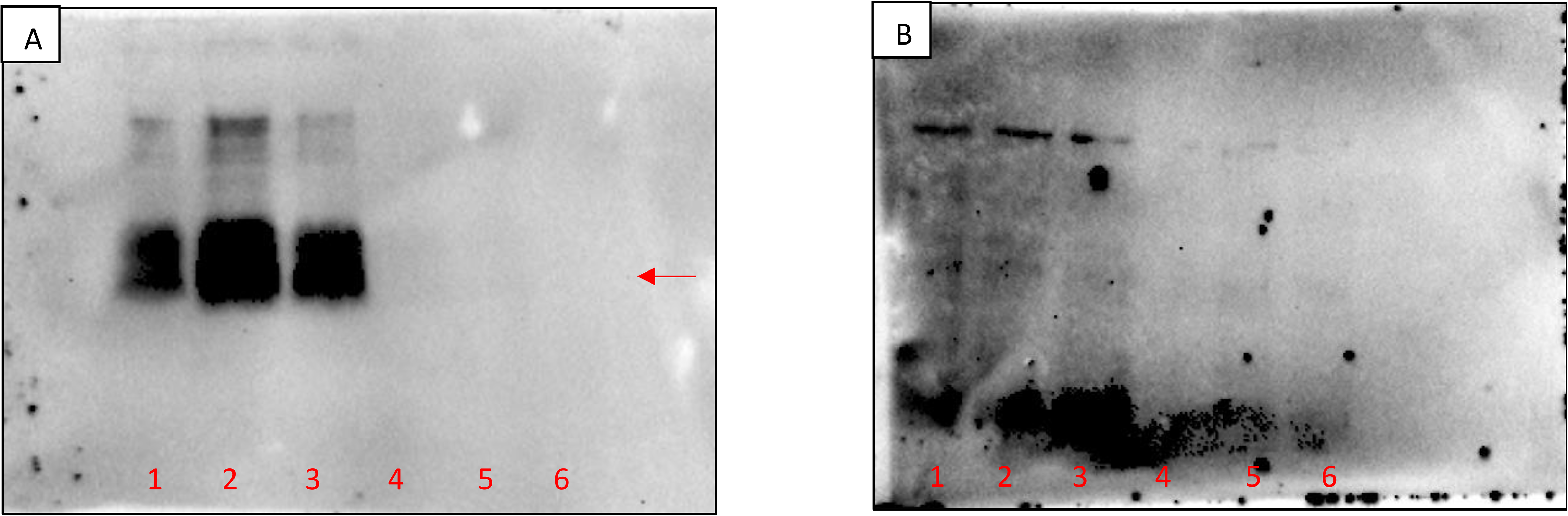
Full phosphorylated tau (pT231) western blots for HEK-293 cells (A) and SH-SY5Y cells (B). Lanes 1-3: non-transfected cells; lanes 4-6: transfected cells.

**Figure S6:**
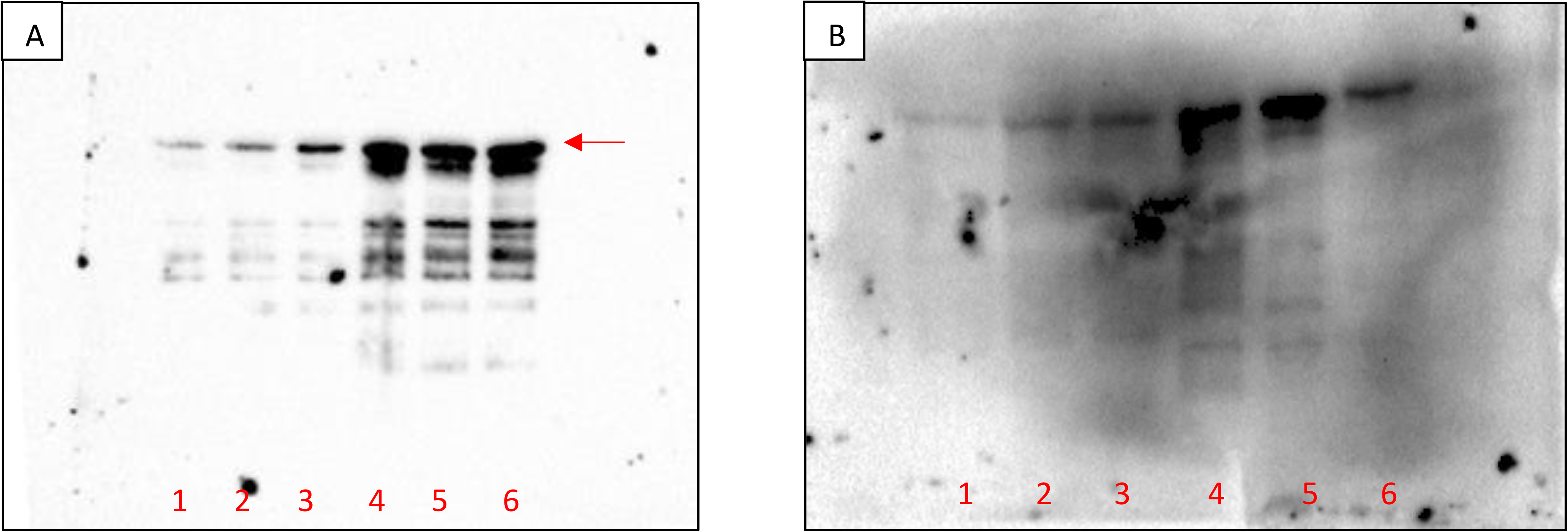
Full hTERT western blots for HEK-293 cells (A) and SH-SY5Y cells (B). Lanes 1-3: non-transfected cells; lanes 4-6: transfected cells.

**Figure S7:**
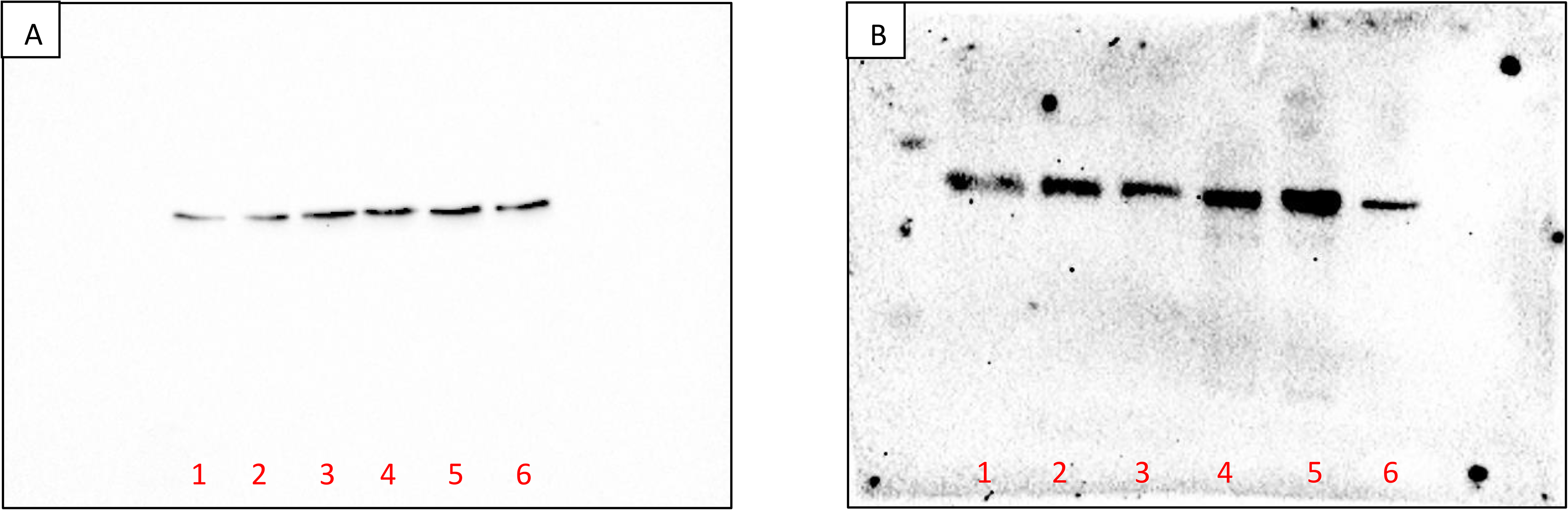
Full phosphorylated TERT (pS824) western blots for HEK-293 cells (A) and SH-SY5Y cells (B). Lanes 1-3: non-transfected cells; lanes 4-6: transfected cells.

**Figure S8:**
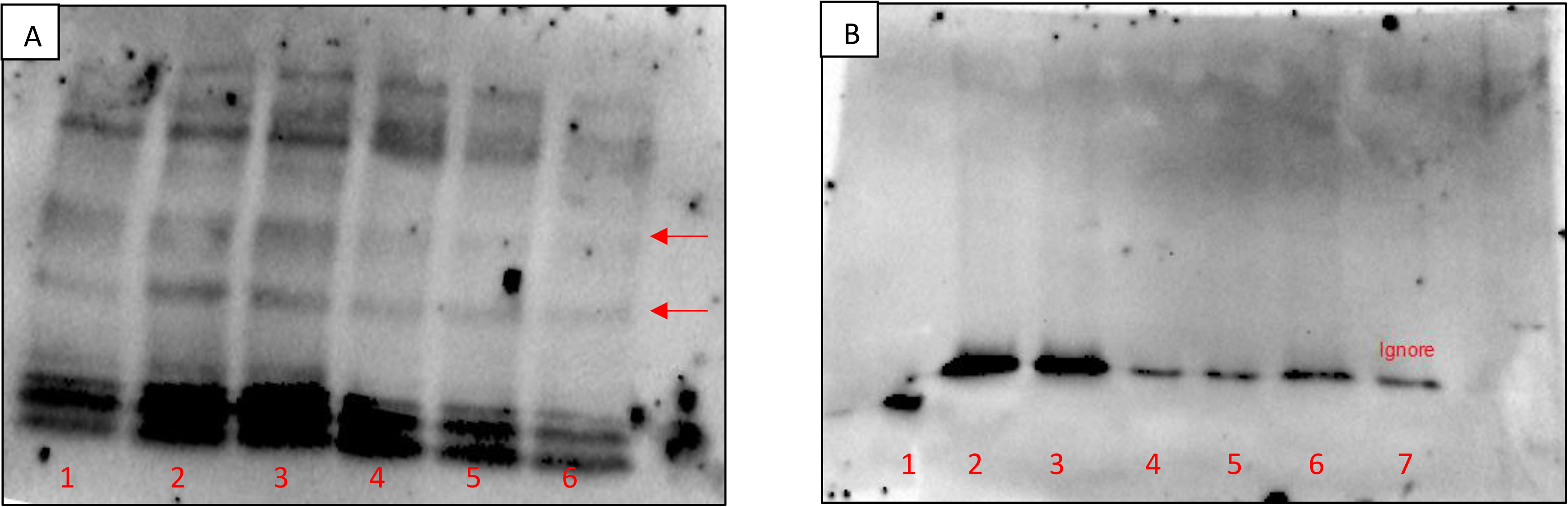
Full PrP^c^ western blots for HEK-293 cells (A) and SH-SY5Y cells (B). Lanes 1-3: non-transfected cells; lanes 4-6: transfected cells; lane 7: repeat of transfected SH-SY5Y cells not included in analysis.

